# Emergence of Canonical Functional Networks from the Structural Connectome

**DOI:** 10.1101/2020.09.16.300384

**Authors:** Xihe Xie, Pablo F. Damasceno, Chang Cai, Srikantan Nagarajan, Ashish Raj

**Affiliations:** Department of Neuroscience, Weill Cornell Medicine, New York, NY 10028; Department of Radiology and Biomedical Imaging, University of California, San Francisco, San Francisco, CA 94143

**Keywords:** structural connectivity, functional networks, graph Laplacian, complex Laplacian

## Abstract

How do functional brain networks emerge from the underlying wiring of the brain? We examine how resting-state functional activation patterns emerge from the underlying connectivity and length of white matter fibers that constitute its “structural connectome”. By introducing realistic signal transmission delays along fiber projections, we obtain a complex-valued graph Laplacian matrix that depends on two parameters: coupling strength and oscillation frequency. This complex Laplacian admits a complex-valued eigen-basis in the frequency domain that is highly tunable and capable of reproducing the spatial patterns of canonical functional networks without requiring any detailed neural activity modeling. Specific canonical functional networks can be predicted using linear superposition of small subsets of complex eigenmodes. Using a novel parameter inference procedure we show that the complex Laplacian outperforms the real-valued Laplacian in predicting functional networks. The complex Laplacian eigenmodes therefore constitute a tunable yet parsimonious substrate on which a rich repertoire of realistic functional patterns can emerge. Although brain activity is governed by highly complex nonlinear processes and dense connections, our work suggests that simple extensions of linear models to the complex domain effectively approximate rich macroscopic spatial patterns observable on BOLD fMRI.

## 1 Introduction

The exploration of structure and function relationships is a fundamental scientific inquiry at all levels of biological organization, and the structure-function relationship of the brain is of immense interest in neuroscience. Attempts at mathematical formulations of neuronal activity began with describing currents traveling through a neuron’s membranes and being charged via ion channels[1]. Recently, the focus of computational models have expanded from small populations of neurons to macroscale brain networks, which are now available via diffusion-weighted and functional magnetic resonance imaging (dMRI and fMRI) [2]. Using computational tractography on dMRI images, detailed whole brain white-matter tracts, and their connectivity can be obtained, to yield the brain’s structural connectivity (SC). Using correlated activation patterns over time in fMRI data reveals functional connectivity (FC) with high spatial resolution. Such high resolution images of the brain also allowed neuroscientists to label the brain according to anatomical or functional regions of interest (ROIs) [3, 4]. Subsequently, efforts in graph-theoretic modeling have emerged as an effective computational tool to study the brain’s SC-FC relationship based on the parcellated brains: ROIs become nodes and connectivity strengths become edges on the graph, while dynamical systems describing neuronal activity are played out on this graph structure [5, 2, 6].

Diverse graph based methods have been employed to relate the brain’s SC to FC. Particularly, perturbations and evolution of the structural and functional networks have been investigated using both graph theoretical statistics [7, 8, 9, 10, 11, 12, 13] as well as network controllability [14, 15]. Structurally informed models use graphical representations of the brain’s connections to couple anatomically connected neuronal assemblies [16, 17], numerical simulations of such neural mass models (NMMs) provides an approximation of the brain’s local and global activities, and are able to achieve moderate correlation between simulated and empirical FC [18, 19, 20, 21, 22]. However, approximations through stochastic simulations are unable to provide a closed form solution and inherits interpretational challenges since dynamics is only obtained from iterative optimizations of high dimensional NMM parameters.

An emergent field of work have suggested low-dimensional processes involving diffusion or random walks on the structural graph as a simple means of simulating FC from SC. These simpler models are equally if not more successful at simulating fMRI FC patterns [23, 24] as well as MEG oscillatory patterns [25, 26] than conventional NMMs. Lastly, these simpler graph diffusion models, which naturally employ the Laplacian of SC, have been generalized to yield spectral graph models whereby Laplacian eigen-spectra were sufficient to reproduce functional patterns of brain activity, using only a few eigenmodes [27, 24, 26]. Thus, a Laplacian matrix representation of a network can be used to find characteristic properties of the network [28], and its eigenmodes (or spectral basis) are the ortho-normal basis that represent particular patterns on the network. Such spectral graph models are computationally attractive due to low-dimensionality and more interpretable analytical solutions.

The SC’s Laplacian eigenmodes are therefore emerging as the substrate on which functional patterns of the brain are thought to be established via almost any reasonable process of network transmission [27, 24, 29], and metrics quantifying structural eigenmode coupling strength to functional patterns were also recently introduced [30]. These works mainly focused on replicating canonical functional networks (CFNs), which are stable large scale circuits made up of functionally distinct ROIs distributed across the cortex that were extracted by clustering a large fMRI dataset [31]. In [31] seven CFNs (these are spatial patterns, not to be confused for the entire network of graph of the connectome) were identified. Hence recent graph modeling work has attempted to address whether these canonical patterns can emerge by only looking at the structural connectivity information of the brain.

Although spectral graph models have been reasonably successful, they leave several important gaps. First, they accommodate only passive spread, hence are incapable of producing oscillating or traveling phenomena, which are critical properties of brain functional activity. Second, they do not incorporate path delays caused by finite axonal conductance speed of activity propagating through brain networks. Third, they are capable of reproducing only deterministic and steady-state features of empirical brain activity, giving a single predicted FC for a given SC. Hence these models cannot easily explain the substantial variability observed amongst individuals, as well as between different recording session of the same individual. This suggests that simplistic spectral graph models will need to be augmented with a set of richer time- or individual-varying features or parameters in order to make them more realistic. Unfortunately, this is a goal that is at variance with the key attraction of these methods - their parsimony and low-dimensionality.

In this study we propose a novel spectral graph approach that is able to produce a far richer range of functional activity and dynamics without compromising on the simplicity and parsimony of the spectral graph model. We hypothesise that the introduction of realistic path delays and axonal conductance speeds can allow graph spectra to display the kinds of pattern-richness observed in real data. Hence we utilize both the SC connectivity strength matrix as measured by white-matter fiber tract density, as well as the distance matrix as measured by the average white-matter fiber tract distance between pairs of ROIs. We show that the additional distance information allows for examining of network dynamics in the complex domain in terms of a novel complex-valued Laplacian. This approach involves only global model parameters, which between them accommodate a rich diversity of spatiotemporal patterns that are capable of closely reproducing the diversity of spatial patterning seen across a large number of healthy subjects. Through this minimalist complex diffusion model, the characteristic patterns of signal spread described by corresponding complex-valued eigen-spectra can be tuned to exhibit activation patterns resembling human CFNs. We show that the complex approach significantly and consistently exceeds the performance of existing works relating real-valued SC Laplacian’s eigen-spectra to measured FC [24, 30, 27, 21]. The introduction of the complex-valued Laplacian and accompanying complex graph diffusion may be an important contribution to the emerging literature on graph models of brain activity, and furthers our understanding of the structure-function relationship in the human brain.

We begin with a general theory of complex graph diffusion incorporating path delays, leading to the emergence of the complex-valued Laplacian. Then we present detailed statistical analysis showing the ability of complex eigenmodes to be tuned by model parameters and reproducing CFNs. We present comparison with the current approach of using real-valued eigenmodes, followed by a detailed Discussion.

## 2 Theory

**Notation.** In our notation, vectors and matrices are represented in **bold**, and scalars by normal font. We denote frequency of a signal, in Hertz (Hz), by symbol *f*, and the corresponding angular frequency as *ω* = 2*πf*. The structural connectivity matrix is denoted by ***C*** = *c_jk_*, consisting of connection strength *cj_k_* between any two pairs of brain regions *j* and *k*.

### 2.1 Network Diffusion of Brain Activity

For an undirected, weighted graph representation of the structural network *c_i,j_*, we model the average neuronal activation rate for the *i*-th region as *x_i_*(*t*):

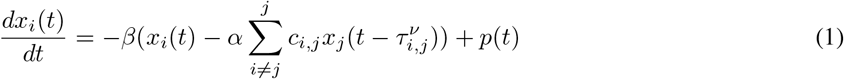

Where we have a mean firing rate equation at the *i*-th region controlled by an inverse of the common characteristic time constant *β*, and input signals from the *j*-th regions connected to region *i* are scaled by the connection strengths from *c_i,j_* and delayed by 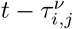. The term 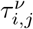 is the delay in seconds obtained from the distance adjacency matrix defined by 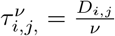, with *ν* representing the conductance speed in the brain’s SC network. The global coupling parameter *α* acts as a controller of weights given to long-range white-matter connections.

The delays between connected brain regions turn into phase shifts in the frequency profiles of the oscillating signals. Thus we obtain the following Fourier transforms from (1): 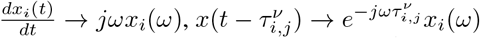, and the oscillatory frequency *ω* = 2*πf*. Lastly, we define a complex connectivity matrix as a function of angular frequency *ω* as 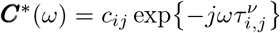. Therefore, a structural connectivity matrix whose nodes are normalized by 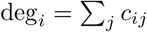 at frequency *ω* can be expressed as:

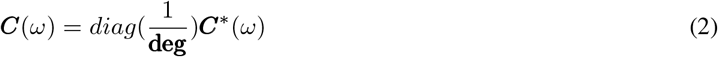

### 2.2 Complex Laplacian Matrix

Our goal is to examine the characteristic patterns of diffusion revealed by the structural network’s normalized Laplacian matrix. Here, we make use of (2) to introduce a complex Laplacian matrix that absorbs the network properties of both the structural connectivity matrix as well as the distance adjacency matrix. By applying the Fourier transforms mentioned above, we obtain a closed-form solution for 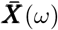:

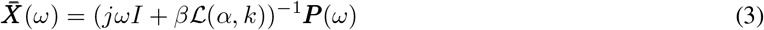

In this closed-form solution, we define a complex Laplacian matrix 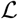 as a function of global coupling *α*. Since frequency *ω* and transmission speed *ν* always occur as a ratio, we define a wave number 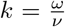. The wave number represents the spatial frequency of any propagating wave, describing the amount of oscillations per unit distance traveled [32]. Then the complex Laplacian matrix 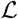 has the form:

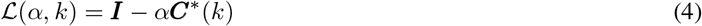

Where ***I*** is the identity matrix and ***C***^*^(*k*) is the complex connectivity matrix as defined above. While (3) indicates that the propagating signals in the network is governed by 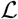, the complex Laplacian of the network describes the characteristic patterns of signal spread in a network, and we can obtain these spatial patterns via the decomposition:

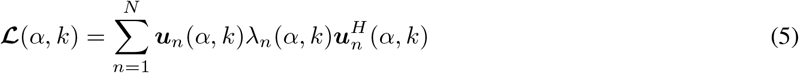

Where λ_*n*_(*α, k*) are the eigenvalues of the complex Laplacian matrix and ***u**_n_*(*α, k*)’s are the complex eigenmodes of the complex Laplacian matrix. Here, the entries of the complex Laplace eigenmodes represent the relative amount of activation in each parcellated brain region as controlled by global coupling and wave number parameters.

**Figure 1:**
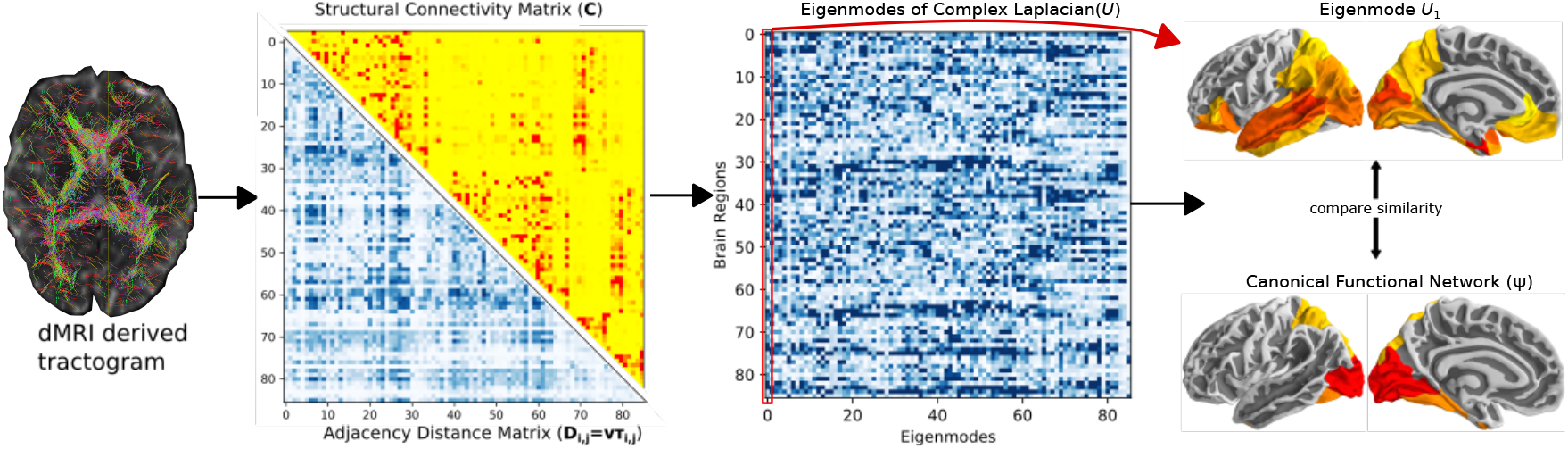
The analysis overview. Structural connectivity matrix (*C*) and distance adjacency matrix (*D*) were extracted from diffusion MRI derived tractograms, to construct the complex Laplacian of the brain’s structural network. An eigen decomposition on the network’s complex Laplacian 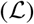 was performed obtain complex structural eigenmodes of the brain (*U*). The spatial similarities were computed between the structural eigenmodes and canonical functional networks in fMRI. Here, as an example, we show brain rendering of the leading eigenmode from the HCP template structural connectome (right column, top) and the canonical visual functional network (right column, bottom).

## 3 Results

### 3.1 Structural connectivity based functional activation patterns

We use the HCP template connectome to demonstrate the wide range of spatial activity patterns achievable by the eigenmodes of the complex Laplacian matrix. The top row of Figure 2 shows three exemplary real-valued structural eigenmodes (*α* = 1) without frequency and transmission speed tuning. Consistent with previous works, we see the Laplace eigenmodes of the human structural connectome display a wide range of cortical activity patterns [27, 24]. As a comparison, we show in the next row complex Laplace eigenmodes with low wave number (*k* = 0.1), representing a network with extremely high transmission speed or near zero delay. In such a low delay network, the complex Laplace eigenmdoes closely resemble the spatial patterns seen in real-valued Laplace eigenmodes where delays are not a factor in the network. We also show two additional examples of complex Laplace eigenmodes with higher wave number values, emphasizing the impact of transmission speed and delays in the structural network of the brain. The combination of coupling strength and wave number global parameters enables a richer diversity of spatial cortical patterns, with left and right hemisphere specific activations around the dorsal-caudal brain regions. Despite the increase in model complexity, our approach allows a feature-rich graph theoretics approach to directly infer resting state functional brain patterns from the structural graph of the brain.

**Figure 2:**
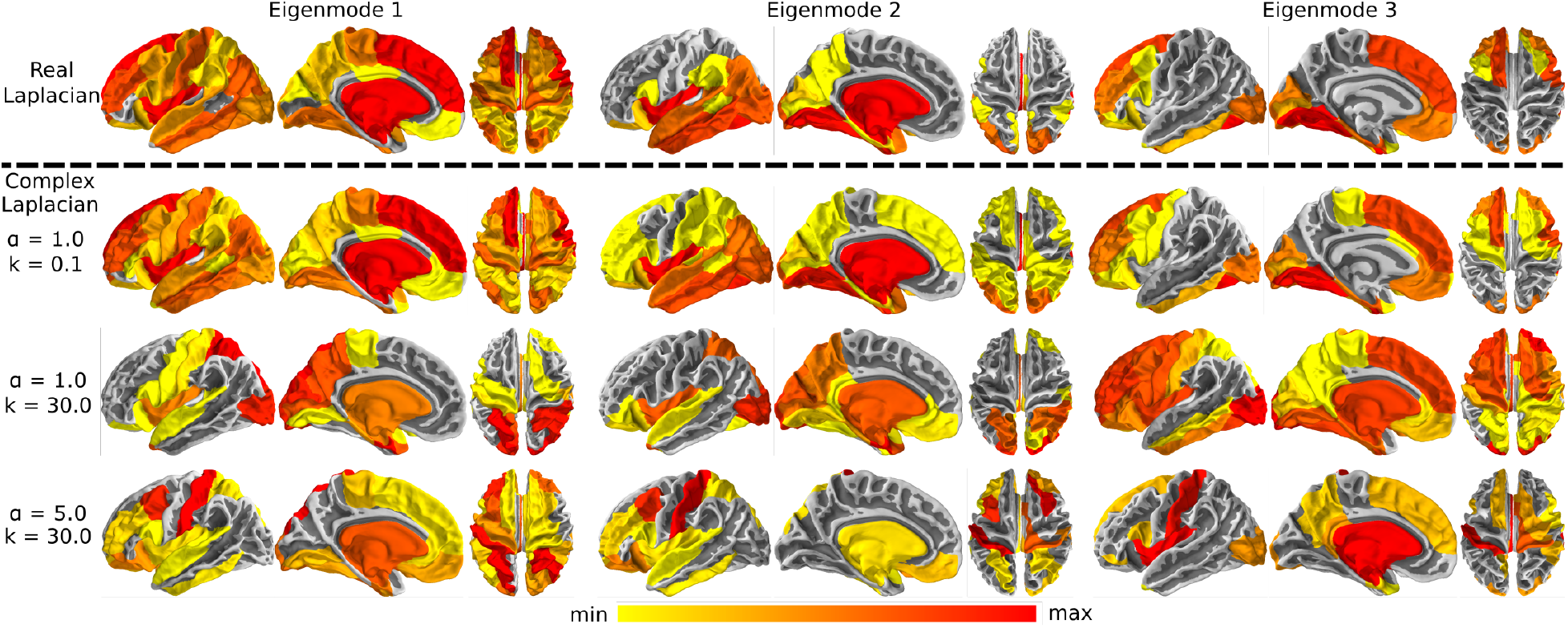
Complex Laplacian eigenmode for different parameter choices. Three representative eigenmodes decomposed from the complex Laplacian with different tuning parameters and three representative eigenmodes decomposed from the real-valued Laplacian without transmission speed and distance delay properties are shown. The top row shows brain renderings of the real Laplacian eigenmodes with coupling strength *α* = 1. Complex Laplacian eigenmodes with high transmission speed approaches extremely small wave number or delays in the network (*α* = 1, *k* = 0.1), closely resembles the real Laplacian eigenmodes (second row). Complex Laplacian eigenmodes with higher wave numbers with parameters (*α* = 1, *k* = 30) and (*α* = 5, *k* = 30) are respectively shown in the third and fourth rows, demonstrating that parameter choice control the spatial distribution of structural eigenmodes.

### 3.2 Eigenmodes of the complex Laplacian resemble CFN activation patterns

We re-assigned the voxel-wise parcellations of the seven CFNs from Yeo et al. [31] to brain regions from the Desikan-Killiany atlas (Figure 3, left column), this re-sampling of the parcellations allow spatial pattern comparisons of equal dimensions against our structural connectomes and Laplace eigenmodes. The middle column of Figure 3 shows best matching complex Laplace eigenmodes after optimization of the global parameters with the HCP template connectome to each CFN. In addition to displaying the best-performing eigenmode in each case, we further ranked the eigenmodes according to their spatial correlation values and displayed the best weighted linear combination of the top 10 complex Laplace eigenmodes on the right column of Figure 3. The spatial correlation values of the best performing eigenmode, and details of cumulative combinations of eigenmodes are reported below and in Figure 6. We observe that CFN patterns emerge when parameters, optimized for each network, are applied to the complex Laplacian. Only a few structural eigenmodes are required to capture a specific functional network.

**Figure 3:**
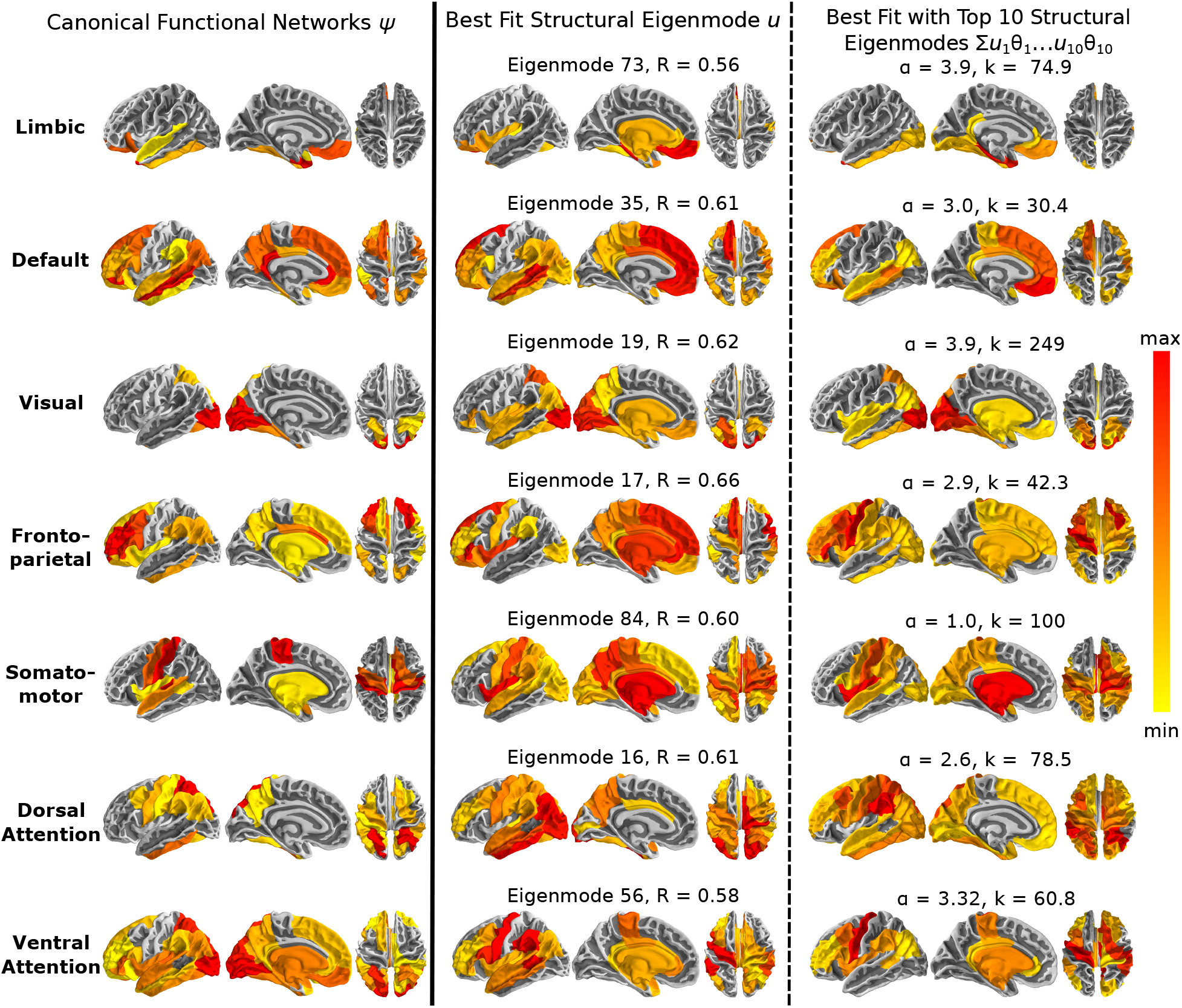
Canonical functional networks reproduced by structural eigenmodes. Brain renderings of the seven canonical functional networks are shown in the left column. Individual structural eigenmodes with the highest spatial correlation to each functional network, after parameter optimization, are shown in the middle column. After ranking all structural eigenmodes by highest spatial correlation, a linear combination of the top ten best performing eigenmodes are shown in the right column. Parameter values producing the best spatial matches to each canonical functional network are listed in the right column and applies to all eigenmodes.

**Figure 4:**
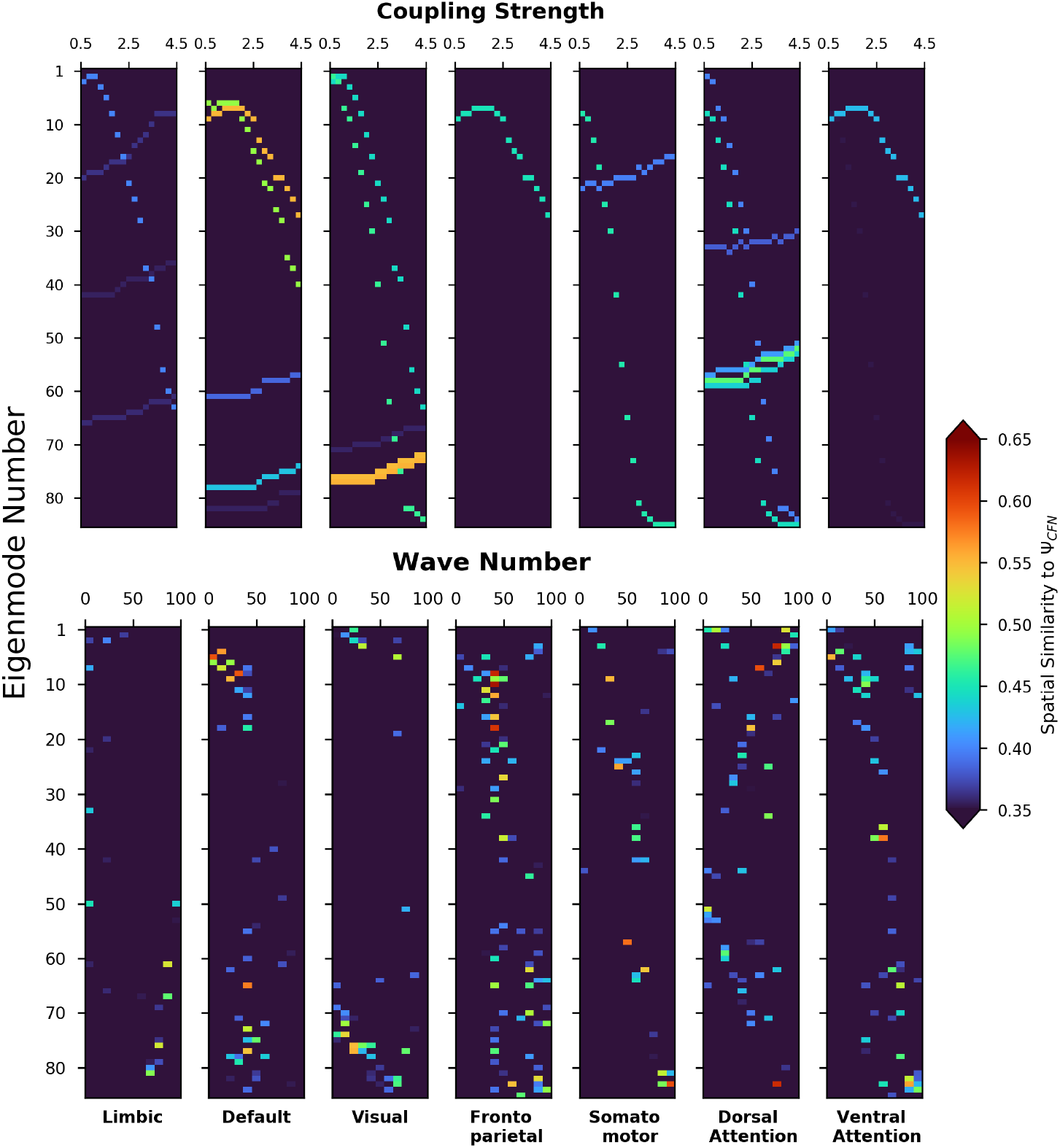
Structural eigenmode spatial similarity to canonical functional networks depends on model parameters. Colors display the spatial correlation values (Spearman’s) of all complex Laplacian eigenmodes across all parameter values with each canonical functional network. Shifts in coupling strength (*α*, top, with wave number held constant at *k* = 10) does not cause a change in peak spatial correlation, but only in the ordering of the eigenmodes. In contrast, however, shifts in wave number (k, bottom), with coupling strength held constant at *α* = 1, leads to changes in eigenmode spatial patterns and spatial correlation to canonical functional networks.

**Figure 5:**
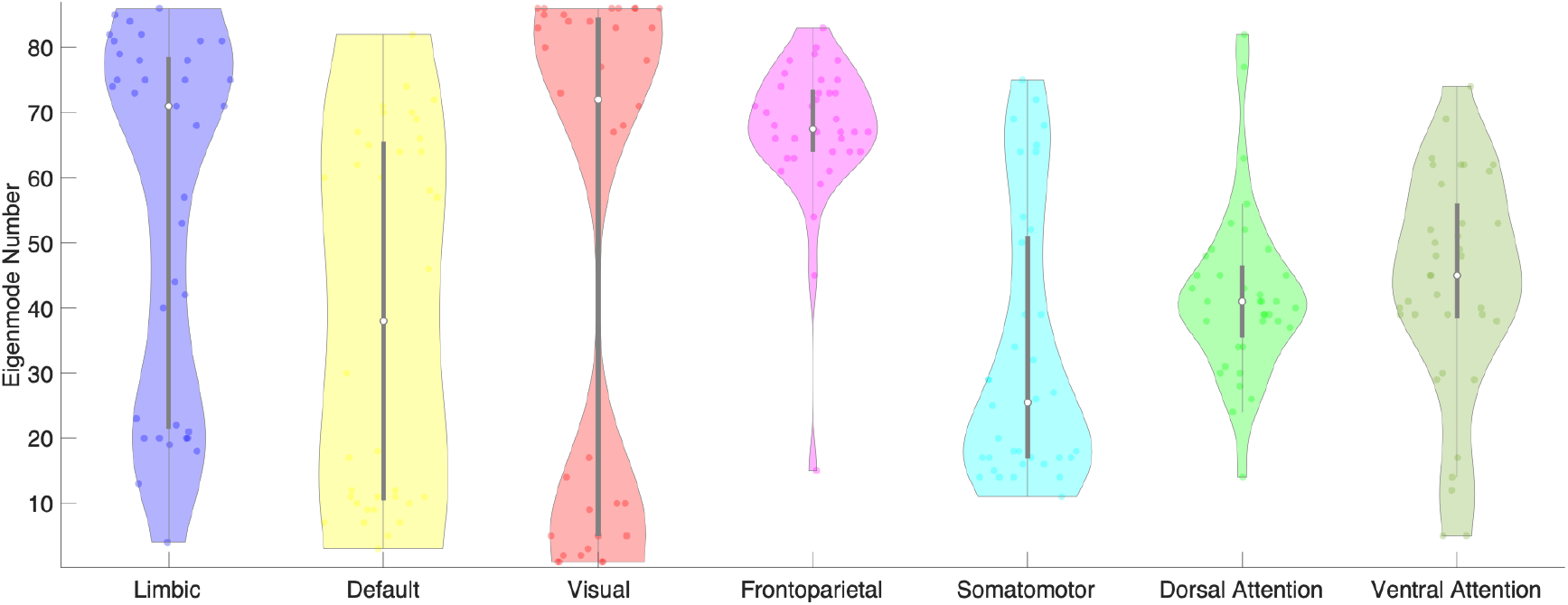
Canonical functional networks have complex Laplacian eigenmode specificity. Each dot on the violin plot corresponds to the best performing eigenmode number. Showing that across all subjects (*n* = 36), canonical functional networks occupies specific structural eigenmodes as the dominant structural basis. Default mode network is the exception as the best performing eigenmode spans across all eigenmodes. On the other hand, the rest of the canonical functional networks cluster to specific eigenmode numbers.

**Figure 6:**
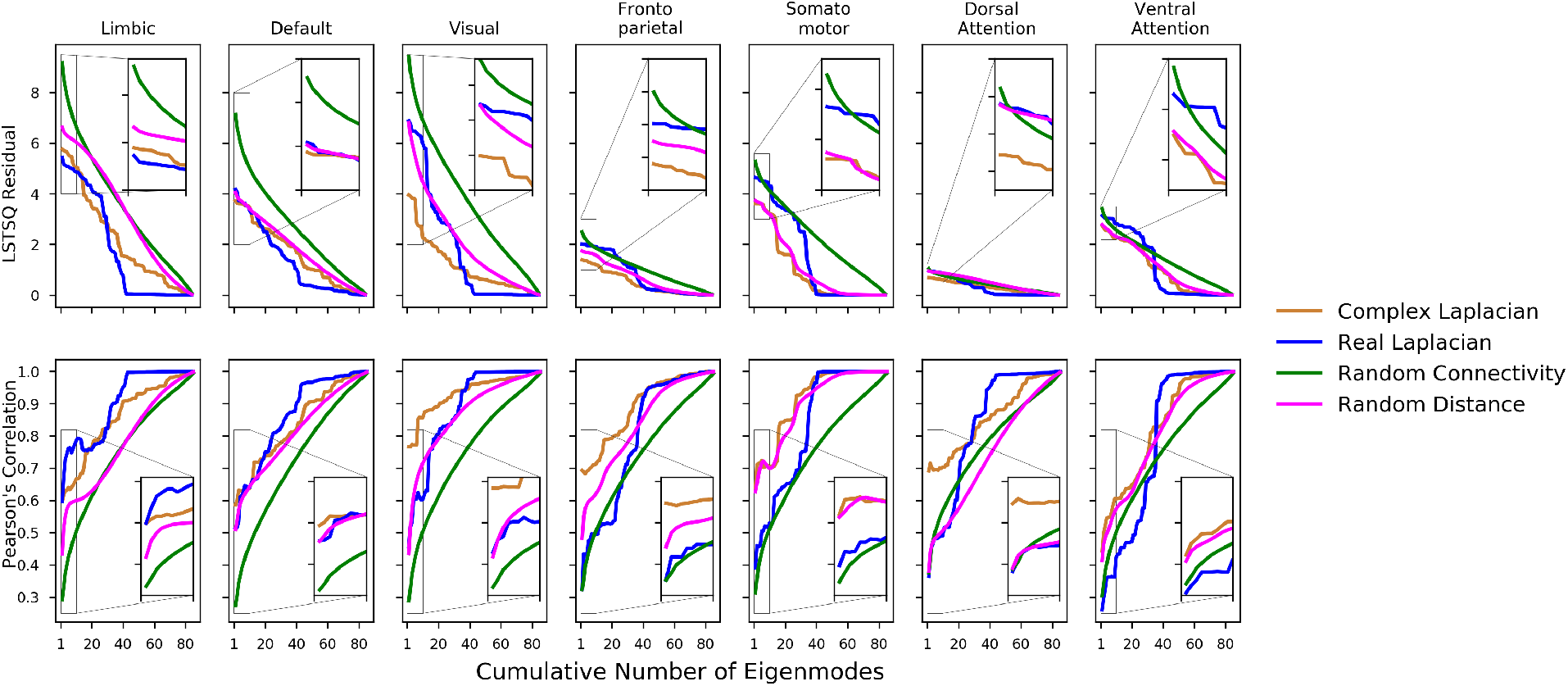
Structural eigenmodes of the HCP template complex Laplacian predict canonical functional networks better than structural eigenmodes of the real Laplacian. For each canonical functional network, we quantified its spatial similarity against linear combinations of structural eigenmodes obtained from various types of Laplacians. The spatial similarities quantified by linear least squares residuals are shown on top, and Pearson’s correlations are shown on the bottom. Overall, accumulation of structural eigenmodes improves the spatial similarity between functional networks and structural eigenmodes. The complex HCP eigenmodes (orange) and real-valued HCP eigenmodes (blue) both outperform eigenmodes decomposed from random connectomes and random distance matrices (green). However, only the complex HCP eigenmodes outperform complex eigenmodes decomposed from the HCP template connectome paired with random distance matrices (magenta).

### 3.3 Parameter tuning of complex Laplacian eigenmodes

To examine the sensitivity of our eigenmodes to our complex Laplacian parameters, we first computed spatial correlation values for the all eigenmodes for each CFN across the entire parameter range. Figure 4 (top) shows the effect of fixing *k* and varying *α*, while bottom row shows the effect of varying *k* at a fixed *α*. At a glance, almost all eigenmodes are capable of resembling any given CFN with the proper choice of tuning parameters, and it is evident that we need to tune both the global coupling strength and wave number for a dominant eigenmode matching a specific CFN to emerge. For any given CFN, we find parameter regimes that recruit multiple eigenmodes while others recruit a single one. This is especially true of the wave number parameter and not so for coupling strength. Furthermore, the best achievable spatial correlation stay consistent as we tweak the global coupling strength, whereas wave number tuning causes shifts in spatial similarity value and eigenmode occupation. And finally, the limbic network has the lowest spatial match and the least a mount of shift in spatial correlation values.

To further examine the tunable parameter’s effects on the leading (best performing) eigenmodes, we show a heat map of the spatial correlation achieved by the dominant eigenmode as we shifted parameter values in Supplementary Figure 1. As expected, global coupling parameter had no effect on dominant eigenmode’s fit while the wave number did. Subsequently, we split the wave number parameter into its two components: transmission velocity and oscillating frequency of signals in the network, showing that those two components equally affect spatial patterns emerging from the complex Laplacian eigenmodes (Supp. Figure 1 bottom row). The spatial correlation patterns of each functional network also implies that there are potentially functional network specific eigenmodes obtainable from the structural complex Laplacian, which will be explored further in the subsequent group level analysis.

On the group level, we found parameter sets that provided the most spatially similar complex Laplace eigenmode for each canonical functional network. The rank of the most spatially similar eigenmodes are summarized in violin plots in Figure 5. With the exception of the default mode network, whose best structural match spans across the range of all eigenmodes, all other canonical functional networks exhibit selectivity towards a specific subset of ranked eigenmodes. The limbic and visual networks, which contains dense connections in the anterior and ventral regions of the brain, prefer to occupy eigenmodes at both low and high ends of the eigen spectrum. On the other hand, the dorsal and ventral attention networks mainly occupy the middle of the eigen-spectrum. The specific occupancy patterns shown here implies there may be a hierarchy to the functional and structural organization of the brain, and the functioning brain minimizes the recruitment of unrelated structural connections when engaged in conscious brain activity.

### 3.4 Complex Laplacian eigenmodes outperform real Laplacian eigenmodes

We created 1000 random realizations of connectivity matrices and their corresponding distance adjacency matrices that share the same sparsity, mean, and standard deviation values as the HCP template connectome values. Comparisons between eigenmodes of the HCP template connectome and randomly generated connectomes are displayed in Figure 6. The Laplace eigenmodes of the brain’s white matter network can be seen as individual subsets of cortical activation patterns that make up the brain’s functional activity. Therefore, spatial match between cumulative combinations of eigenmodes to each canonical functional network were computed in addition to just the leading eigenmode.

Overall, the HCP complex Laplacian’s best-performing eigenmodes achieved higher spatial correlation and lower residuals than other variants of Laplace eigenmodes in 6 out of the 7 CFNs (left-most point on each curve). As more individual eigenmodes are linearly combined, all variants show a steady improvement in spatial similarity, with the fully random variant using the most number of eigenmodes to achieve a high spatial match, suggesting the fully random eigenmodes are the least informative. On the other hand, the complex eigenmodes from randomized distance Laplacians (magenta) consistently performs better than fully random complex eigenmodes (green) but lacks the structural distance information to compete with complex Laplace eigenmodes constructed with HCP template connectivity and distance adjacency matrix. The spatial similarity reported in Figure 6 is Pearson’s correlation due to its smoothness, we show the same quantification with Spearman’s correlation in Supplementary Figure 2, which is more appropriate for discrete samples, but its more volatile due to its nonlinear ordering of samples.

Spatial similarities from random variants of complex Laplace eigenmodes were normalized into a Z-score distribution for construction of 95% confidence intervals and comparisons against HCP connectome variants. Comparing only the leading eigenmodes without cumulative combinations, Complex Laplacian eigenmodes significantly outperforms random connectivity eigenmodes for all functional network comparisons (*P* < 0.05), but only significantly outperforms the randomized distance eigenmodes for the limbic, visual, frontoparietal, and dorsal attention networks. On the other hand, the real-valued Laplace eigenmodes does not significantly outperform eigenmodes from fully random connectivity profiles for all functional networks. The *P*-values for both Pearson’s and Spearman’s metrics are shown in Supplementary Tables S1 and S2.

### 3.5 Group level eigenmode analysis

Figure 7 shows a violin plot of the best spatial correlation achieved by each subject’s complex Laplacian in orange, real Laplacian in blue, and random distance adjacency matrix paired with the HCP connectome in magenta. Consistent with our HCP template connectome analysis, the complex Laplacian eigenmodes outperforms both the real Laplacian eigenmodes and randomized distance complex Laplacian eigenmodes. Our complex Laplacian framework includes the additional distance and delay information in the brain networks compared to conventional real Laplacian eigenmodes, therefore we generated complex Laplacien eigenmodes from HCP structural connectivity paired with random distance adjacency matrices, which as a comparative degree of freedom. Paired T-tests were performed for all CFNs, the complex Laplacian eigenmodes outperformed real Laplcian eigenmodes at the group level for all networks except the limbic network (*p* = 0.64). On the other hand, significantly higher spatial similarity was achieved by complex Laplacian eigenmodes for all networks except the dorsal attention network (*p* = 0.12) when comparing against the random distance group results.

**Figure 7:**
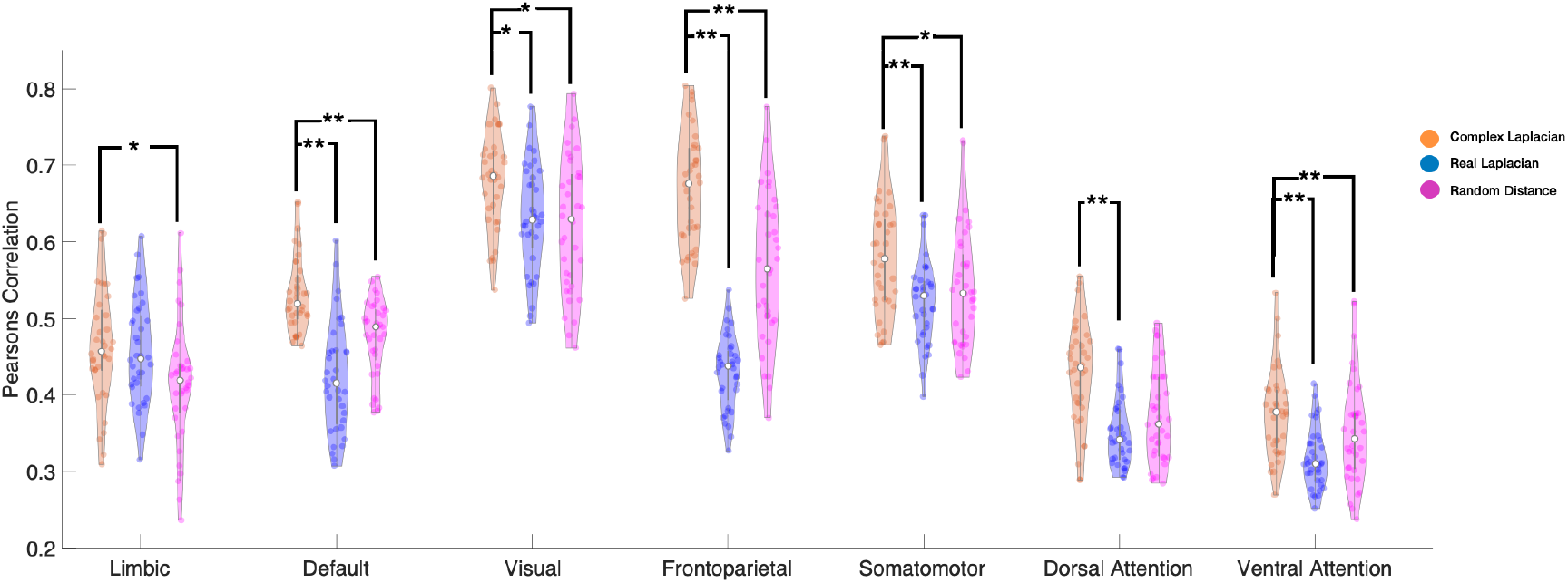
Complex Laplacian outperforms real Laplacian in recapitulating canonical functional networks with individual structural connectomes. Violin plot showing that on a group level (each dot correspond to one subject, n = 36), the best performing structural eigenmodes of the complex Laplacian (orange) outperforms the corresponding structural eigenmode from the real Laplacian (blue) and random distance complex Laplacian (magenta). Paired T-test results of complex Laplacian against either real Laplacian or random distance complex Laplacian shows the complex Laplacians eigenmodes achieving significantly higher spatial similarity on the group level (P-values shown as * < 0.5, ** < 0.01).

## 4 Discussion

In this study we have proposed a complex graph Laplacian framework that demonstrates an ability to capture functional connectivity patterns, while maintaining parsimony and low-dimensionality of spectral graph models. The model involves only two global and biophysically meaningful parameters, one controlling the speed of activity propagation, and the other controlling coupling strength between remote populations of neurons connected via axonal projections. We presented detailed statistical analysis of the resulting complex-value Laplacian eigenmodes, focusing on their ability to predict the spatial patterns observed on seven CFNs that are well established in functional neuroimaging. The implications of our main contributions are discussed below, with additional context and relevance to current literature.

### 4.1 A simple yet feature-rich graph theoretic approach

We derived a simple model of network diffusion of activity which takes into account the path delays introduced by realistic axonal conductance speeds and fiber lengths, and showed that at the first order the behavior of the model can be captured within a complex Laplacian, on which a complex-valued graph diffusion process is enacted. Using this definition of the complex Laplacian we demonstrated that its eigenmodes constitute a sparse basis that is capable of reproducing the characteristic spatial patterns of empirical resting state functional activity given by the 7 CFNs.

### 4.2 Higher predictive power than existing graph models

We showed that the complex Laplacian outperforms the existing models that use the eigenmodes of real-valued Laplacian. These results are far better than can be expected by chance, as indicated by the significance values of our results with respect to large simulations with Laplacians calculated from random connectomes. Thus, future graph models can benefit from the enhanced predictive power of the proposed complex Laplacian approach, which in the cases we have tested highly significantly improves performance(see Figure 5). Our work can therefore find direct applicability in many clinical and neuroscientific contexts where predicting functional patterns from structure is important [33, 34], particularly in cases of epilepsy [35], stroke [36, 37], and neurodegeneration [38].

### 4.3 Complex eigenmodes accommodate a diversity of spatial patterns

One of the most intriguing aspects of our study is the demonstration that almost all (complex) eigenmodes are capable of resembling any given CFN, with the proper choice of tuning parameters. As observed from Figure 4, certain parameter regimes recruit multiple eigenmodes while others recruit a single one; however with the right selection of the two model parameters, it is possible to “steer” the eigenmodes in such a manner that a small number of them can reproduce any CFN. This not only denotes the strength of our approach, we believe it points to an essential characteristic of real brain activity, which is thought to accommodate a large repertoire of microstates and their concomitant spatial patterns. This rich repertoire was shown above to be capable of being engaged by our parsimonious graph model, which may point to the possibility that complex behavior may be achievable by simple and parsimonious mechanisms, and may not require the kinds of high-dimensional and non-linear oscillatory models that have held sway in the field of neural modeling [21, 19]. Our work also supports the idea that macroscopic neurophysiological data on a graph can be sufficiently modeled with linear metrics, and nonlinear methods may not be required for problem of such scale [39, 40].

### 4.4 Rich repertoire is tunable with two biophysical parameters

In our model, the brain can access any configuration of spatial patterns seen in real resting state functional networks by tuning only two of its global and biophysically meaningful parameters: coupling strength and wave number. Our current work indicates that physical distances and the transmission rate of oscillatory activity in combination with coupling strength is sufficient in generating various canonical functional brain patterns. This demonstration in an analytical model, that a rich repertoire of states is accessible to the brain by tuning biophysical processes, has not previously been reported to our knowledge. The present computational study is not intended to explore the neural mechanisms that might control these parameters. Nevertheless, modern neuroscience provides several potential mechanisms.

Coupling strength *α* is a direct scaling of white-matter excitatory long range connections between neural populations in the brain. Phase and amplitude coupling of oscillatory processes in the brain is evidently important for the formation of coherent wide-band frequency profiles of brain recordings and processing of information [41, 42, 43, 44, 45]. Parameterization of coupling strength between distant brain regions via the connectome is ubiquitous in connectivity based models of BOLD fMRI [46, 45, 21, 23] and electroencephalography activity [47, 22, 48]. Furthermore, pathological FC patterns as a result of disconnections in the brain can be reproduced with decrease in coupling strength [49, 50].

The other key tunable parameter in our model, wave number *k*, is the ratio between the oscillatory frequency and transmission velocity of a propagating signal, describing the amount of oscillations per unit distance traveled by any signal spreading throughout the brain’s structural network. While transmission speed of signals between brain regions is often overlooked in brain modeling efforts, its importance is emphasized by the biology of the central nervous system. Neuronal spike arrival timing at the cellular level and coherent oscillatory activity at the network level are carefully managed by synaptic strengths as well as axonal myelination, respectively [51, 52]. Further, wave number can be controlled not just by conductance speed, but also by the operative frequency of oscillations *ω*. From the deep literature on wide-band frequency response of brain recordings, it is already known that different functional networks of resting state BOLD data are preferentially encapsulated by different higher-frequency bands via phase- and amplitude-coupling [44, 45]. Hence it is plausible that wave number tuning may be achieved biologically via either dynamic conductance speed or dynamic control of frequency bands.

### 4.5 Relationship to existing studies

Recent graph models involving eigen spectra of the adjacency or Laplacian matrices of the structural connectome have greatly contributed to our understanding of how the brain’s structural wiring gives rise to its functional patterns of activity (REF). Although these models have very attractive features of parsimony and low-dimensionality, they suffer from being feature poor and an inability to make stronger predictions about functional networks.

Such models mapping between structural and functional patterns of the human brain have typically assumed that SC and FC are not independent entities, and that relationship between the two cannot simply be explained by a direct mapping [21]. In addition to connection strength between regions, metrics such as anatomical distances [53], shortest path lengths [54], diffusion properties [27, 36], and structural graph degree [55] were also found to contribute to the brain’s observed functional patterns. Higher-order walks on graphs have also been quite successful; typically these methods involve a series expansion of the graph adjacency or Laplacian matrices [56, 57]. The diffusion and series expansion methods are themselves closely related [29], and almost all harmonic-based approaches may be interpreted as special cases of each other, as demonstrated elegantly in recent studies [58, 59]. The wealth of studies elucidating how the observed function originate from the underlying structural network provided a strong motivation for our approach, which extracts functional patterns from the informative complex graph Laplacian that incorporates both the connection strengths as well as the anatomical distances of the structural network.

In contrast to spectral graph models, inferring functional connectivity from biophysiological models of neuronal populations have been a specialty of dynamic causal models (DCMs). Such generative models have emerged as powerful tools mainly to infer effective (directional) connectivity for smaller networks [60, 61, 62, 63, 64], or dynamic functional connectivity [65, 66]. While the goal of DCMs is similar to our proposed model that makes model inferences about FC, the two frameworks are different in terms of approach and dimensionality. DCMs examine the second order covariances of brain activity, and it is only recent works with spectral and regression DCM models have expanded the model coverage to the whole-brain scale and the potential to incorporate SC data [67, 68, 69]. However, these models rely on formulation of local neural masses to derive dynamical behavior, which are then used to generate effective or dynamic connectivity through simulations. By avoiding large-scale simulations of neuronal activity, in our proposed framework we not only allowed canonical functional patterns to emerge directly from a complex Laplacian matrix, we have also created a model with only two global parameters. Most DCM models have many more degrees of freedom compared to our work because of their parameterization for different interactions within and between brain regions. In contrast to some of more recent spectral DCM parameterizations, additionally, our global parameters reflecting the brain’s anatomical connection density and distances traveled between connections continue to have clear biophysical interpretability.

Frequency-band specific magnetoencephalography (MEG) resting-state networks have been successfully modeled with a combination of delayed NMMs and eigenmodes of the structural network [25], suggesting delayed interactions in a brain’s network give rise to functional patterns constrained by structural eigenmodes. In our recent work, we expanded upon eigenmdoes of SC matrices by integrating time delays in the brain with SC to create a complex Laplacian matrix in the Fourier domain [26]. Using the eigen-spectra of the complex Laplacian matrix, we found specific subsets of complex eigenmodes that contributed to specific cortical alpha and beta wave patterns. The findings in the current article expands upon these time-delayed eigenmodes to find subsets of eigenmodes predictive of canonical functional networks derived from resting state fMRI. Our theorized framework provides two global parameters that act on the structural connectome and its corresponding distance adjacency matrix to control coupling strength and delays in the network. These findings supports other works suggesting there is a possible organizational hierarchy, or gradients of topographical organization that spatially constraints cortical function [70, 71, 72, 73, 74]. Margulies et al. proposed that so-called “principal gradients”, which may be interpreted as the Laplacian eigenmodes of the FC matrix, serve as the core organizing axis of cerebral cortex, spanning from unimodal sensorimotor to integrative transmodal areas [70]. The complex eigenmodes proposed here may therefore be considered as the structural analog of Margulies’ principal gradients. Similarly, we found that the unimodal sensorimotor networks at one end of the principal gradient, which accounts for the most variance in connectivity, achieved the highest spatial correlations. On the other hand, transmodal networks on the opposite end of the axis, needed much more cumulatively combined structural eigenmodes to achieve high spatial similarity.

Atasoy et al. previously modeled the same resting-state canonical functional networks used here with real-valued Laplacian eigenmodes as structural substrates on which a mean field neural model dictated cortical dynamics [24]. While the model dimensionalities between the two studies are vastly different, we show that in the absence of a neural dynamical system, the addition of time lag in the network allowed canonical functional networks to emerge from just structural substrates. Furthermore, we believe incorporating time lags in our structural connectivity of the brain to create complex Laplacian matrices is an informative but unexplored alternative to regular Laplacian normalizations of brain networks. Particularly, the complex connectivity matrix in Fourier domain allows exploration of oscillatory frequency and phase shifts between brain regions as a property of the network, potentially presenting an opportunity in utilizing complex structural eigenmodes to integrate SC for explaining imaginary coherence patterns in MEG and EEG.

### 4.6 Limitations

The current results are limited by data resolution. Tractograms obtained from diffusion weighted images are approximations of the brain’s axonal white-matter connections. We recognize that tractography, paired with anatomical parcellation of brain regions, does fail to appreciate the finer structures in the brain, especially the more refined connections and nuclei in the brain stem as well as close neighbor connections. Despite the coarse parcellation and rough approximations of white matter architecture, our proposed approach utilizes a spatial embedding of the brain’s connectomics information and is extendable to finer parcellations.

Our theorized model relies on an averaged approximation of fiber distances between ROIs, and we assumed a global parameter to account for conductance speed in the brain. In reality, the amount of myelination and synaptic strength varies greatly in the brain. However, our approximations were enough in recapitulating canonical functional networks in the human brain, while benefiting from a low dimensional and interpretable model. It is also worth nothing that the canonical functional networks used in this work were obtained from data-driven clustering of fMRI activity, and is far from a comprehensive representation of the brain’s functional patterns. While our work can be extended to finer functional parcellations, we sought to avoid overlap between canonical functional networks by using the 7 networks parcellation. For example, the dorsal and ventral attention networks are found to overlap with the salience network [75], and task activated fMRI patterns revealed regions that are positively and negatively associated with attention and default networks [76].

## 5 Conclusions

In conclusion, we show that the spatial embedding of the brain’s connections in a structural connectome is a rich substrate, on which we can derive intrinsic functional patterns of the brain with a simple network diffusion approach. We show that Laplace eigenbasis in the complex frequency domain outperforms conventional eigenbasis of the graph Laplacian in capturing spatial patterns of canonical functional networks. We recognize the complex nonlinear activities and dense connections present in the brain, but our work suggests that we can continue to extend simpler linear modeling approaches to approximate what we observe with macroscopic imaging techniques such as BOLD fMRI and diffusion weighted imaging.

## 6 Methods

### 6.1 Structural Connectivity Network Computation

We constructed structural connectivity networks according to the Desikan-Killiany atlas where the brain images were parcellated into 68 cortical regions and 18 subcortical regions as available in the FreeSurfer software [77, 3]. We first obtained openly available diffusion MRI data from the MGH-USC Human Connectome Project to create an average template connectome [78]. Additionally, we obtained individual structural connectivity networks from 36 subjects’ diffusion MRI data. Specifically, *Bedpostx* was used to determine the orientation of brain fibers in conjunction with *FLIRT*, as implemented in the *FSL* software [79]. Tractography was performed using *probtrackx2* to determine the elements of the adjacency matrix. We initiated 4000 streamlines from each seed voxel corresponding to a cortical or subcortical gray matter structure and tracked how many of these streamlines reached a target gray matter structure. The weighted connection between the two structures *c_i,j_* was defined as the number of streamlines initiated by voxels in region *i* that reach any voxel within region *j*, normalized by the sum of the source and target region volumes. This normalization prevents large brain regions from having extremely high connectivity due to having initiated or received many streamline seeds. Afterwards, connection strengths are averaged between both directions (*c_i,j_* and *c_i,j_*) to form undirected edges. Additionally, to determine the geographic location of an edge, the top 95% of non-zero voxels by streamline count were computed for both edge directions, the consensus edge was defined as the union between both post-threshold sets.

### 6.2 Canonical Functional Networks

We chose the 7 CFN parcellations mapped by Yeo et al. [31] as the functional spatial patterns most frequently visited by the human brain. The brain parcellations were created from fMRI recordings of 1000 young, healthy English speaking adults at rest with eyes open. A clustering algorithm was used to parcellate and identify consistently coupled voxels within the brain volume. The results revealed a coarse parcellation of seven networks: **Ψ**_*CFN*_ = {limbic, default, visual, frontoparietal, somatomotor, ventral attention, dorsal attention}.

The CFN parcellation was co-registered to brain regions of interest in the gyral based Desikan-Killany atlas [3] to match the dimensionality of our complex Laplacian structural eigenmodes. Then spatial activation maps of each canonical network was produced by normalizing the number of voxels per brain region belonging to a specific CFN by the total number of voxels in the brain region of interest (Fig 1). Both the functional networks and the Desikan-Killiany atlas are openly available for download from Freesurfer [80] (http://surfer.nmr.mgh.harvard.edu/).

### 6.3 Global Parameter Optimization for Individual Structural Eigenmodes

To ensure that we obtained a globally optimal set of parameters *α, k* that provided a complex Laplacian eigenmode ***u**_n_* which is the most similar to the spatial pattern of each of the seven **Ψ**_*CFN*_, we performed an optimization of the cost function: *f*(*α, k, n*) = 1 – *corr*(**Ψ**_*CFN*_, ***u**_n_*(*α, k*)) to determine the optimal eigenmode, coupling, and wavenumber for each canonical functional network. We used the “basin-hopping” global optimization technique on this cost function, a robust technique for non-convex cost functions [81]. This algorithm is able to escape from local minima in the parameter space by accepting and “hopping” to new parameters even if they increase the cost function. The algorithm will accept iterations that decrease the cost function evaluation with a probability of 1, but only accept iterations that do not decrease cost function with a probability of exp(Δ(*f*)/*T*), where Δ(*f*) is the change in the cost function across successive iterations, and T is a constantly decreasing “temperature” term. Larger T indicates that the algorithm is more willing to accept jumps in cost function evaluation. We initiated the optimization procedure from ten different initial parameter values and selected the best result out of all initialization runs.

### 6.4 Similarity Analysis Between Canonical Functional Networks and Cumulative Linear Combination of Structural Eigenmodes

Here, we examine whether structural eigenmodes can form a linear basis for activation patterns for canonical functional networks and examine if a cumulative combination of structural eigenmodes improves the spatial similarity with CFN’s when compared to individual structural eigenmodes. For each CFN, we first ordered the eigenmodes based on their individual similarity after global parameter optimization using procedures described in the previous section. For each CFN, we then computed similarity of the optimal linear weighting of sorted individual structural eigenmodes **u**_*l*_ with **Ψ**_*CFN*_ by cumulatively adding structural eigenmodes ordered by their similarity. We minimized the *L*_2_ – norm of 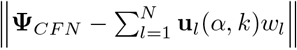, to obtain the optimal weights *w_l_* and a quantification of spatial patterns obtained by the best cumulative set of eigenmodes.

Spatial similarity of cumulative eigenmodes with CFNs were then computed using both Pearson’s (Figure 6) and Spearman’s correlations (Figure S2). While Spearman’s correlation was appropriate for non-continuous correlative comparisons, its non-linearity due to sorting of values was evident in volatile changes of spatial similarity, and Pearson’s correlation provided more stable results.

We repeated the above analysis for both the conventional real-valued Laplacian without frequency and transmission speed tuning, as well as complex Laplacians obtained from randomized connectivity matrices. For random connectivity matrices, we constructed 1000 realizations of random connectivity and distance matrices to allow us to compare and quantify the performance of the brain’s structural eigenmodes against eigenmodes of randomized graphs. The random matrices were constructed with the same sparsity as the HCP template connectome, and the elements of the random matrices were assigned by randomly sampling from a distribution that’s representative of the mean and variance of the HCP template connectome and distance matrices.

## 7 Code & Data Availability

Intermediate light-weight data and code that support the findings of this study are available from the GitHub repository at https://github.com/axiezai/complex_laplacian. The code used to produce basic figures can be run as interactive Jupyter notebooks after installing the computing environment from https://zenodo.org/record/3532497 [82], instructions for downloading and setting up the computing requirements are documented in the README file.

## 8 Supplementary Material

**Figure S1:**
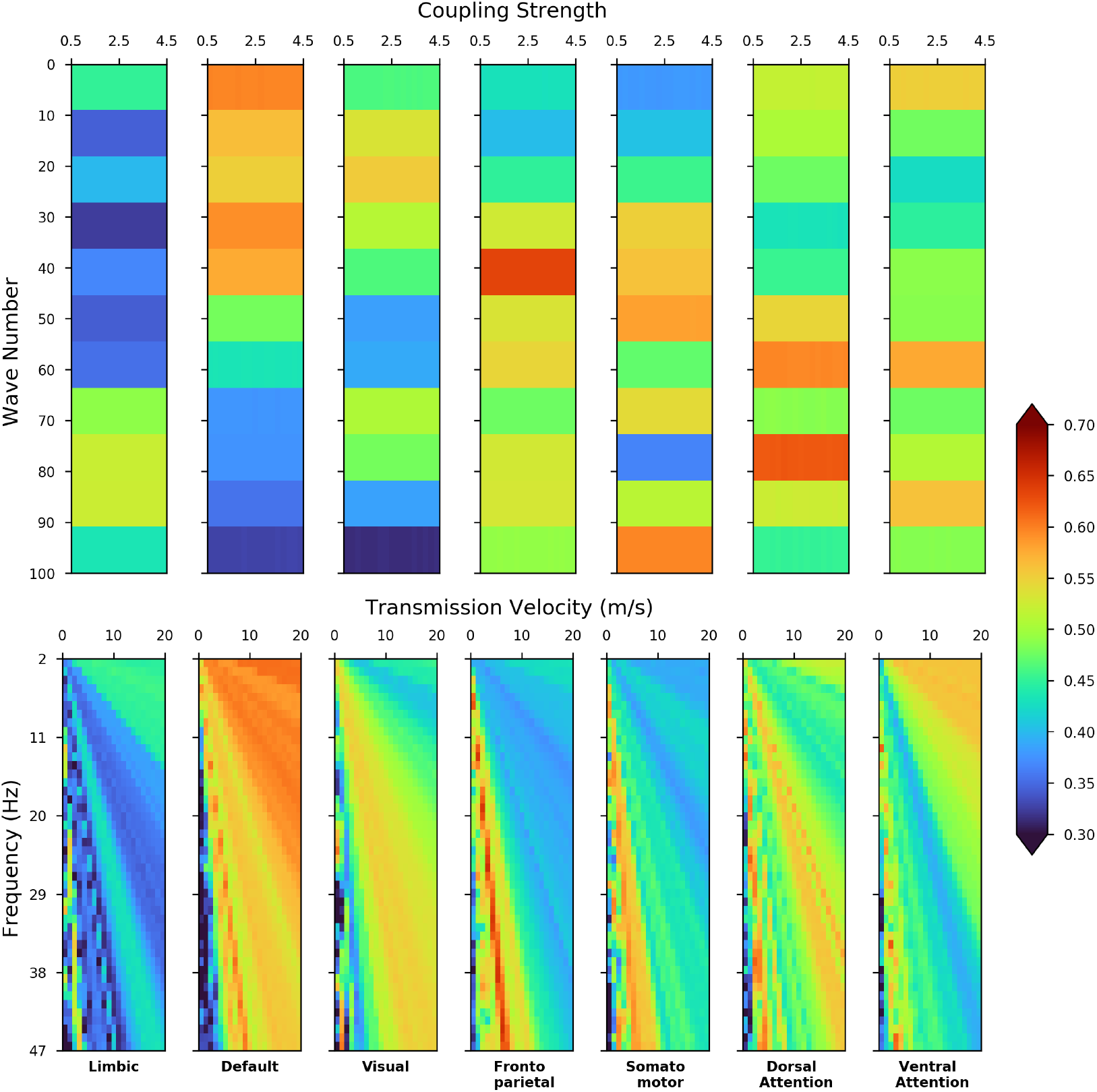
Parameter dependency of structural complex Laplace eigenmodes. Heat-maps displaying best achievable spatial correlation values (Spearman’s) by a single eigenmode across all parameter values for each canonical functional network. Shifts in coupling strength (top) does not cause a change in peak spatial correlation. The bottom row shows the transmission speed and oscillating frequency of signals in the network dictates the cortical activation patterns in the brain, shifts in wave number parameters while holding global coupling constant changes the best achievable spatial similarity to each canonical network.

**Table S1:** *P*-values table from random connectome comparisons of leading eigenmodes. Z-score distributions of spatial correlation (Pearson’s) were created from 1000 sets of complex Laplace eigenmodes of 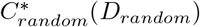 and 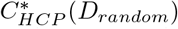 random connectomes. For all canonical networks’ similarity comparisons, a 95% confidence interval of the Z-scores distributions were obtained and used to compute the *P*-values shown in the tables.

| | $C_{HCP}^*$ vs. $C_{random}^*$<br>($D_{random}$) | $C_{HCP}$ vs. $C_{random}^*$<br>($D_{random}$) | $C_{HCP}^*$ vs. $C_{HCP}^*$<br>($D_{random}$) | $C_{HCP}$ vs. $C_{HCP}^*$<br>($D_{random}$) |
| --- | --- | --- | --- | --- |
| **Limbic** | 1.504e-06 | 1.466e-06 | 1.355e-03 | 1.328e-03 |
| **Default** | 2.015e-07 | 5.512e-05 | 1.119e-01 | 4.804e-01 |
| **Visual** | 1.122e-13 | 4.967e-03 | 9.580e-06 | 3.928e-01 |
| **Frontoparietal** | 2.263e-06 | 4.993e-01 | 2.856e-03 | 1.789e-02 |
| **Somatomotor** | 5.882e-06 | 1.459e-01 | 3.088e-01 | 9.173e-23 |
| **Dorsal Attention** | 2.412e-04 | 4.522e-01 | 6.038e-06 | 4.101e-01 |
| **Ventral Attention** | 2.700e-02 | 2.705e-01 | 6.175e-02 | 6.202e-16 |

**Figure S2:**
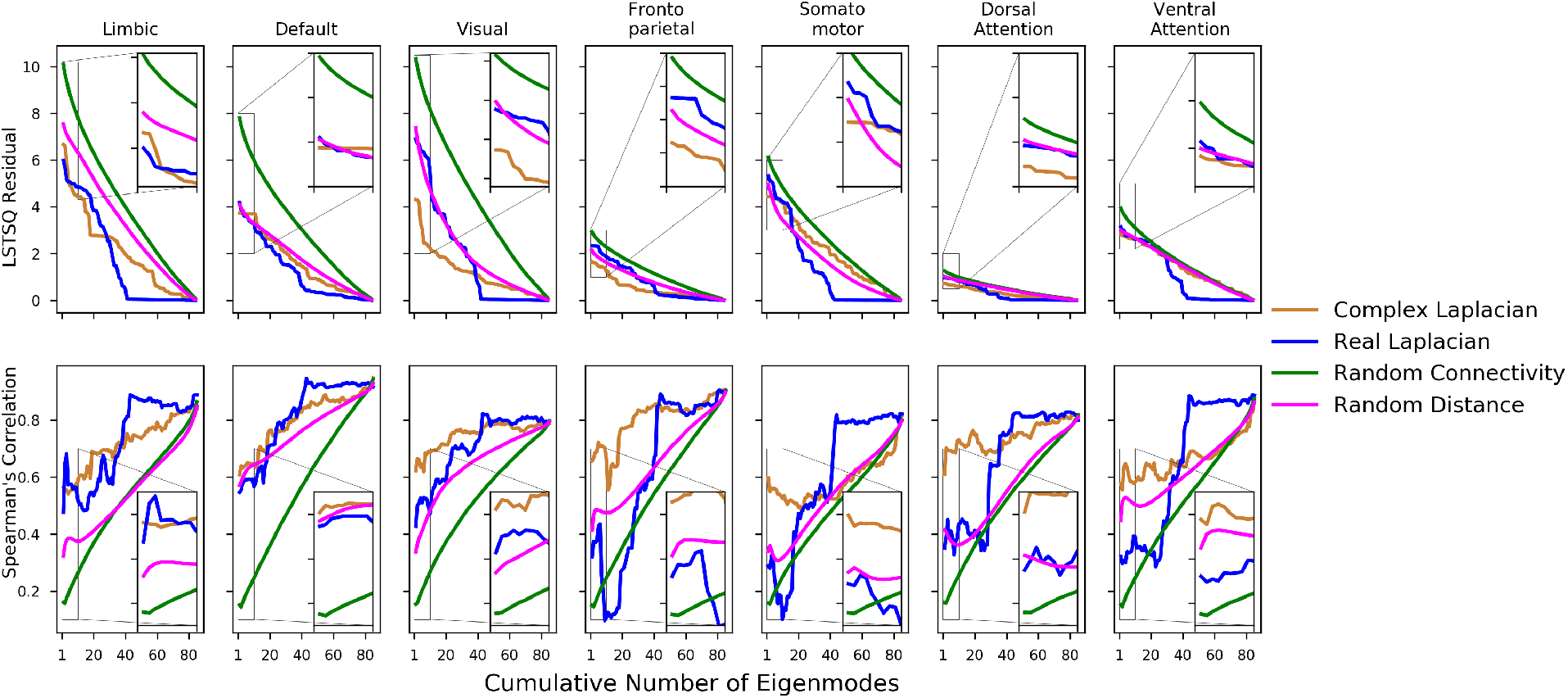
Spatial pattern similarity of HCP template connnectome complex Laplacian structural eigenmodes to canonical functional networks quantified with Spearman’s correlation. The same analysis performed in Figure 2 but Pearson’s correlation was replaced with Spearman’s correlation for discrete samples for the bottom row, and the top row shows linear least square residuals. Despite the more inconsistent increasing trend in spatial pattern similarity due to ranking of discrete samples, the complex Laplacian eigenmodes are able to outperform the real-valued Laplacian eigenmodes (blue), complex eigenmodes from random structural connectome with random distance matrix (green), as well as complex eigenmodes from HCP template connectome with random distance matrix.

**Table S2:** *P*-values table from random connectome comparisons of leading eigenmodes. This table is produced the same way as Table S1, but the Z-score distributions were computed from Spearman’s correlation

| | $C_{HCP}^*$ vs. $C_{random}^*$<br>($D_{random}$) | $C_{HCP}$ vs. $C_{random}^*$<br>($D_{random}$) | $C_{HCP}^*$ vs. $C_{HCP}^*$<br>($D_{random}$) | $C_{HCP}$ vs. $C_{HCP}^*$<br>($D_{random}$) |
| --- | --- | --- | --- | --- |
| **Limbic** | 4.423e-06 | 2.448e-04 | 4.593e-07 | 8.516e-04 |
| **Default** | 1.404e-07 | 4.469e-06 | 2.872e-01 | 3.573e-01 |
| **Visual** | 7.601e-08 | 1.091e-03 | 1.312e-05 | 9.155e-02 |
| **Frontoparietal** | 3.181e-09 | 2.455e-02 | 4.866e-05 | 6.873e-02 |
| **Somatomotor** | 4.160e-07 | 7.129e-02 | 5.856e-07 | 1.616e-01 |
| **Dorsal Attention** | 5.086e-07 | 1.955e-02 | 7.333e-03 | 2.072e-01 |
| **Ventral Attention** | 1.312e-06 | 3.077e-02 | 4.396e-02 | 4.676e-02 |

## References

[1] A. L. Hodgkin and A. F. Huxley. “A quantitative description of membrane current and its application to conduction and excitation in nerve”. In: The Journal of Physiology 117.4 (Aug. 1952), pp. 500–544. ISSN: 0022-3751. URL: https://www.ncbi.nlm.nih.gov/pmc/articles/PMC1392413/.

[2] Ed Bullmore and Olaf Sporns. “Complex brain networks: graph theoretical analysis of structural and functional systems”. en. In: Nature Reviews Neuroscience 10.3 (Mar. 2009), pp. 186–198. ISSN: 1471-0048. DOI: 10.1038/nrn2575. URL: https://www.nature.com/articles/nrn2575.

[3] Rahul S. Desikan et al. “An automated labeling system for subdividing the human cerebral cortex on MRI scans into gyral based regions of interest”. In: NeuroImage 31.3 (July 2006), pp. 968–980. ISSN: 10538119. DOI: 10.1016/j.neuroimage.2006.01.021. URL: https://linkinghub.elsevier.com/retrieve/pii/S1053811906000437.

[4] R. Cameron Craddock et al. “A whole brain fMRI atlas generated via spatially constrained spectral clustering”. In: Human brain mapping 33.8 (Aug. 2012). ISSN: 1065-9471. DOI: 10.1002/hbm.21333. URL: https://www.ncbi.nlm.nih.gov/pmc/articles/PMC3838923/.

[5] Danielle S. Bassett and Edward T. Bullmore. “Human brain networks in health and disease”. eng. In: Current Opinion in Neurology 22.4 (Aug. 2009), pp. 340–347. ISSN: 1473-6551. DOI: 10.1097/WC0.0b013e32832d93dd.

[6] Miao Cao et al. “Topological organization of the human brain functional connectome across the lifespan”. en. In: Developmental Cognitive Neuroscience 7 (Jan. 2014), pp. 76–93. ISSN: 1878-9293. DOI: 10.1016/j.dcn.2013.11.004. URL:http://www.sciencedirect.com/science/article/pii/S1878929313000960.

[7] Nivedita Chatterjee and Sitabhra Sinha. “Understanding the mind of a worm: hierarchical network structure underlying nervous system function in C. elegans”. In: Progress in Brain Research 168 (2008), pp. 145–153. ISSN: 0079-6123. DOI: 10.1016/S0079-6123(07)68012-1.

[8] Danielle Smith Bassett and Ed Bullmore. “Small-World Brain Networks”. In: The Neuroscientist 12.6 (Dec. 1, 2006). Publisher: SAGE Publications Inc STM, pp. 512–523. ISSN: 1073-8584. DOI: 10.1177/1073858406293182. URL: https://doi.org/10.1177/1073858406293182.

[9] Y. He, Z. Chen, and A. Evans. “Structural Insights into Aberrant Topological Patterns of Large-Scale Cortical Networks in Alzheimer’s Disease”. In: Journal of Neuroscience 28.18 (2008). ISBN: 1529-2401 (Electronic)\n0270-6474 (Linking), pp. 4756–4766. ISSN: 0270-6474. DOI: 10.1523/JNEUROSCI.0141-08.2008. URL: http://www.jneurosci.org/cgi/doi/10.1523/JNEUROSCI.0141-08.2008.

[10] R. L. Buckner. “Molecular, Structural, and Functional Characterization of Alzheimer’s Disease: Evidence for a Relationship between Default Activity, Amyloid, and Memory”. In: Journal of Neuroscience 25.34 (2005). ISBN: 0270-6474, pp. 7709–7717. ISSN: 0270-6474. DOI: 10.1523/JNEUROSCI.2177-05.2005. URL: http://www.jneurosci.org/cgi/doi/10.1523/JNEUROSCI.2177-05.2005.

[11] N Brunel and N Brunel. “Dynamics of sparsely connected networls of excitatory and inhibitory neurons”. In: Computational Neuroscience 8 (2000). ISBN: 0929-5313, pp. 183–208. ISSN: 09284257. DOI: 10.1016/S0928-4257(00)01084-6.

[12] Nicolas Brunel and Xiao Jing Wang. “Effects of neuromodulation in a cortical network model of object working memory dominated by recurrent inhibition”. In: Journal of Computational Neuroscience 11.1 (2001). ISBN: 0929-5313, pp. 63–85. ISSN: 09295313. DOI: 10.1023/A:1011204814320. arXiv: gr-qc/9809069v1.

[13] Laura E. Suárez et al. “Linking Structure and Function in Macroscale Brain Networks”. In: Trends in Cognitive Sciences 24.4 (Apr. 1, 2020), pp. 302–315. ISSN: 1364-6613. DOI: 10.1016/j.tics.2020.01.008. URL: http://www.sciencedirect.com/science/article/pii/S1364661320300267.

[14] Shi Gu et al. “Controllability of structural brain networks”. In: Nature Communications 6 (2015). Publisher: Nature Publishing Group, p. 8414. ISSN: 2041-1723. DOI: 10.1038/ncomms9414. URL: http://www.nature.com/doifinder/10.1038/ncomms9414.

[15] Sarah Feldt Muldoon et al. “Stimulation-Based Control of Dynamic Brain Networks”. In: PLoS computational biology 12.9 (2016). arXiv: 1505.02402 ISBN: 1553-734x, e1005076. doi:10.1371/journal. pcbi.1005076. ISSN: 00189286. DOI: 10.1109/TAC.2015.2437520. URL:http://arxiv.org/abs/1601.00987.

[16] H R Wilson and J D Cowan. “Excitatory and inhibitory interactions in localized populations of model neurons.” In: Biophysical journal 12.1 (1972). ISBN: 0006-3495, pp. 1–24. ISSN: 0006-3495. DOI: 10.1016/S0006-3495(72)86068-5. URL: http://www.sciencedirect.com/science/article/pii/S0006349572860685.

[17] Sami El Boustani and Alain Destexhe. “A master equation formalism for macroscopic modeling of asynchronous irregular activity states.” In: Neural computation 21.1 (2009). ISBN: 0899-7667, pp. 46–100. ISSN: 0899-7667. DOI: 10.1162/neco.2009.02-08-710. URL: http://www.ncbi.nlm.nih.gov/pubmed/19210171.

[18] Paul L. Nunez. “The brain wave equation: a model for the EEG”. In: Mathematical Biosciences 21.3 (1974). ISBN: 1600255564, pp. 279–297. ISSN: 00255564. DOI: 10.1016/0025-5564(74)90020-0.

[19] V. K. Jirsa and H. Haken. “A derivation of a macroscopic field theory of the brain from the quasi-microscopic neural dynamics”. In: Physica D: Nonlinear Phenomena 99.4 (Jan. 1, 1997), pp. 503–526. ISSN: 0167-2789. DOI: 10.1016/S0167-2789(96)00166-2. URL: http://www.sciencedirect.com/science/article/pii/S0167278996001662.

[20] P. A. Valdes et al. “Nonlinear EEG analysis based on a neural mass model”. In: Biological Cybernetics 81.5 (Nov. 5, 1999). Publisher: Springer-Verlag ISBN: 0340-1200 (Print) 0340-1200 (Linking), pp. 415–424. ISSN: 03401200. DOI: 10.1007/s004220050572. URL: http://link.springer.com/10.1007/s004220050572.

[21] C J Honey et al. “Predicting human resting-state functional connectivity from structural connectivity.” In: Proceedings of the National Academy of Sciences of the United States of America 106.6 (2009). ISBN: 0027-8424, pp. 2035–40. ISSN: 1091-6490. DOI: 10.1073/pnas.0811168106. URL: http://www.ncbi.nlm.nih.gov/pubmed/19188601.

[22] A. Spiegler and V. Jirsa. “Systematic approximations of neural fields through networks of neural masses in the virtual brain”. In: NeuroImage 83 (March 2013). ISBN: 1053-8119, pp. 704–725. ISSN: 10538119. DOI: 10.1016/j.neuroimage.2013.06.018.

[23] Farras Abdelnour, Henning U. Voss, and Ashish Raj. “Network diffusion accurately models the relationship between structural and functional brain connectivity networks”. In: NeuroImage 90 (2014). Publisher: Elsevier Inc. ISBN: 1095-9572 (Electronic)\r1053-8119 (Linking), pp. 335–347. ISSN: 10538119. DOI: 10.1016/j.neuroimage.2013.12.039. arXiv: NIHMS150003. URL: http://dx.doi.org/10.1016/j.neuroimage.2013.12.039.

[24] Selen Atasoy, Isaac Donnelly, and Joel Pearson. “Human brain networks function in connectome-specific harmonic waves”. In: Nature Communications 7.1 (Apr. 2016), p. 10340. ISSN: 2041-1723. DOI: 10.1038/ncomms10340. URL:http://www.nature.com/articles/ncomms10340.

[25] Prejaas Tewarie et al. “How do spatially distinct frequency specific MEG networks emerge from one underlying structural connectome? The role of the structural eigenmodes”. In: NeuroImage 186 (Feb. 1, 2019), pp. 211–220. ISSN: 1053-8119. DOI: 10.1016/j.neuroimage.2018.10.079. URL: http://www.sciencedirect.com/science/article/pii/S1053811918320603.

[26] Ashish Raj et al. “Spectral graph theory of brain oscillations”. In: Human Brain Mapping n/a.n/a (), pp. 1–19. DOI: 10.1002/hbm.24991. eprint: https://onlinelibrary.wiley.com/doi/pdf/10.1002/hbm.24991. URL: https://onlinelibrary.wiley.com/doi/abs/10.1002/hbm.24991.

[27] Farras Abdelnour et al. “Functional brain connectivity is predictable from anatomic network’s Laplacian eigen-structure”. In: NeuroImage 172 (2018), pp. 728–739. ISSN: 10959572. DOI: 10.1016/j.neuroimage.2018.02.016.

[28] Ian Stewart. “Holes and hot spots”. In: Nature 401.6756 (Oct. 1999), pp. 863–865. ISSN: 0028-0836. DOI: 10.1038/44730. URL:http://www.nature.com/articles/44730.

[29] P. A. Robinson et al. “Eigenmodes of brain activity: Neural field theory predictions and comparison with experiment”. In: NeuroImage 142 (2016), pp. 79–98. ISSN: 10959572. DOI: 10.1016/j.neuroimage.2016.04.050.

[30] Maria Giulia Preti and Dimitri Van De Ville. “Decoupling of brain function from structure reveals regional behavioral specialization in humans”. In: Nature Communications 10.1 (2019). arXiv: 1905.07813. ISSN: 20411723. DOI: 10.1038/s41467-019-12765-7. URL:https://doi.org/10.1038/s41467-019-12765-7.

[31] B T Thomas Yeo et al. “The organization of the human cerebral cortex estimated by intrinsic functional connectivity.” In: Journal of neurophysiology 106.3 (Sept. 2011), pp. 1125–65. ISSN: 1522-1598. DOI: 10.1152/jn.00338.2011. URL: http://www.ncbi.nlm.nih.gov/pubmed/21653723 http://www.pubmedcentral.nih.gov/articlerender.fcgi?artid=PMC3174820.

[32] A.P. French. Vibrations and Waves. M.I.T. introductory physics series. Taylor & Francis, 1971. ISBN: 9780748744473. URL: https://books.google.com/books?id=RqE26vDmd5wC.

[33] Tianzi Jiang. “Brainnetome: a new - ome to understand the brain and its disorders”. In: NeuroImage 80 (Oct. 15, 2013), pp. 263–272. ISSN: 1095-9572. DOI: 10.1016/j.neuroimage.2013.04.002.

[34] Alex Fornito, Andrew Zalesky, and Michael Breakspear. “The connectomics of brain disorders”. In: Nature Reviews. Neuroscience 16.3 (Mar. 2015), pp. 159–172. ISSN: 1471-0048. DOI: 10.1038/nrn3901.

[35] Ana C. Coan et al. “Frequent seizures are associated with a network of gray matter atrophy in temporal lobe epilepsy with or without hippocampal sclerosis”. In: PloS One 9.1 (2014), e85843. ISSN: 1932-6203. DOI: 10.1371/journal.pone.0085843.

[36] A. Kuceyeski et al. “The application of a mathematical model linking structural and functional connectomes in severe brain injury”. In: NeuroImage: Clinical 11 (2016). Publisher: The Authors, pp. 635–647. ISSN: 22131582. DOI: 10.1016/j.nicl.2016.04.006. URL:http://dx.doi.org/10.1016/j.nicl.2016.04.006.

[37] Anne K. Rehme and Christian Grefkes. “Cerebral network disorders after stroke: evidence from imaging-based connectivity analyses of active and resting brain states in humans”. In: The Journal of Physiology 591.1 (Jan. 1, 2013), pp. 17–31. ISSN: 1469-7793. DOI: 10.1113/jphysiol.2012.243469.

[38] J. Zimmermann et al. “Differentiation of Alzheimer’s disease based on local and global parameters in personalized Virtual Brain models”. In: NeuroImage: Clinical 19 (2018). ISBN: 8135449542, pp. 240–251. ISSN: 22131582. DOI: 10.1016/j.nicl.2018.04.017. URL: https://linkinghub.elsevier.com/retrieve/pii/S2213158218301268.

[39] Jaroslav Hlinka et al. “Functional connectivity in resting-state fMRI: is linear correlation sufficient?” In: NeuroImage 54.3 (Feb. 1, 2011), pp. 2218–2225. ISSN: 1095-9572. DOI: 10.1016/j.neuroimage.2010.08.042.

[40] D. Hartman et al. “The role of nonlinearity in computing graph-theoretical properties of resting-state functional magnetic resonance imaging brain networks”. In: Chaos (Woodbury, N.Y.) 21.1 (Mar. 2011), p. 013119. ISSN: 1089-7682. DOI: 10.1063/1.3553181.

[41] Pascal Fries. “A mechanism for cognitive dynamics: neuronal communication through neuronal coherence”. In: Trends in Cognitive Sciences 9.10 (Oct. 1, 2005), pp. 474–480. ISSN: 1364-6613. DOI: 10.1016/j.tics.2005.08.011. URL: http://www.sciencedirect.com/science/article/pii/S1364661305002421.

[42] Alfons Schnitzler and Joachim Gross. “Normal and pathological oscillatory communication in the brain”. In: Nature Reviews Neuroscience 6.4 (Apr. 2005). Number: 4 Publisher: Nature Publishing Group, pp. 285–296. ISSN: 1471-0048. DOI: 10.1038/nrn1650. URL: https://www.nature.com/articles/nrn1650.

[43] Francisco Varela et al. “The brainweb: Phase synchronization and large-scale integration”. In: Nature Reviews Neuroscience 2.4 (Apr. 2001). Number: 4 Publisher: Nature Publishing Group, pp. 229–239. ISSN: 1471-0048. DOI: 10.1038/35067550. URL:https://www.nature.com/articles/35067550.

[44] A. Ghosh et al. “Cortical network dynamics with time delays reveals functional connectivity in the resting brain”. In: Cognitive Neurodynamics 2.2 (Apr. 23, 2008), p. 115. ISSN: 1871-4099. DOI: 10.1007/s11571-008-9044-2. URL: https://doi.org/10.1007/s11571-008-9044-2.

[45] G. Deco et al. “Key role of coupling, delay, and noise in resting brain fluctuations”. In: Proceedings of the National Academy of Sciences 106.25 (2009). ISBN: 0027-8424, pp. 10302–10307. ISSN: 0027-8424. DOI: 10.1073/pnas.0901831106. arXiv: Keyroleofcoupling, delay, andnoiseinrestingbrainfluctuations. (2009). ProceedingsoftheNationalAcademyofSciences, 106(29), 12207âĂŞ12208. doi: 10.1073/pnas.0906701106. URL:http://www.pnas.org/cgi/doi/10.1073/pnas.0901831106.

[46] Christopher J Honey et al. “Network structure of cerebral cortex shapes functional connectivity on multiple time scales.” In: Proceedings of the National Academy of Sciences 104.24 (2007). ISBN: 0027-8424 (Print)\n0027-8424 (Linking), pp. 10240–10245. ISSN: 0027-8424. DOI: 10.1073/pnas.0701519104. URL: http://www.pnas.org/cgi/doi/10.1073/pnas.0701519104 papers3://publication/doi/10.1073/pnas.0701519104.

[47] Anandamohan Ghosh et al. “Noise during rest enables the exploration of the brain’s dynamic repertoire”. In: PLoS Computational Biology 4.10 (2008). ISBN: 1553-7358. ISSN: 1553734X. DOI: 10.1371/journal.pcbi.1000196.

[48] Olivier David and Karl J. Friston. “A neural mass model for MEG/EEG: Coupling and neuronal dynamics”. In: NeuroImage 20.3 (2003). ISBN: 1053-8119 (Print)\r1053-8119 (Linking), pp. 1743–1755. ISSN: 10538119. DOI: 10.1016/j.neuroimage.2003.07.015.

[49] Joana Cabral et al. “Modeling the outcome of structural disconnection on resting-state functional connectivity”. In: NeuroImage 62.3 (Sept. 1, 2012), pp. 1342–1353. ISSN: 1053-8119. DOI: 10.1016/j.neuroimage.2012.06.007. URL: http://www.sciencedirect.com/science/article/pii/S1053811912005848.

[50] V. K. Jirsa et al. “Towards the virtual brain: Network modeling of the intact and the damaged brain”. In: Archives Italiennes de Biologie 148.3 (2010). ISBN: 0003-9829 (Print)\n0003-9829 (Linking), pp. 189–205. ISSN:00039829. DOI: 10.4449/AIB.V148I3.1223.

[51] I Lorena Arancibia-Cárcamo et al. “Node of Ranvier length as a potential regulator of myelinated axon conduction speed”. In: eLife 6 (Jan. 28, 2017). Ed. by Klaus-Armin Nave. Publisher: eLife Sciences Publications, Ltd, e23329. ISSN: 2050-084X. DOI: 10.7554/eLife.23329. URL: https://doi.org/10.7554/eLife.23329.

[52] R. Douglas Fields. “A new mechanism of nervous system plasticity: activity-dependent myelination”. In: Nature Reviews Neuroscience 16.12 (Dec. 2015). Number: 12 Publisher: Nature Publishing Group, pp. 756–767. ISSN: 1471-0048. DOI: 10.1038/nrn4023. URL: https://www.nature.com/articles/nrn4023.

[53] Aaron F. Alexander-Bloch et al. “The Anatomical Distance of Functional Connections Predicts Brain Network Topology in Health and Schizophrenia”. In: Cerebral Cortex (New York, NY) 23.1 (Jan. 2013), pp. 127–138. ISSN: 1047-3211. DOI: 10.1093/cercor/bhr388. URL: https://www.ncbi.nlm.nih.gov/pmc/articles/PMC3513955/.

[54] Joaquín Goñi et al. “Resting-brain functional connectivity predicted by analytic measures of network communication”. In: Proceedings of the National Academy of Sciences 111.2 (Jan. 14, 2014). Publisher: National Academy of Sciences Section: Biological Sciences, pp. 833–838. ISSN: 0027-8424, 1091-6490. DOI: 10.1073/pnas.1315529111. URL: https://www.pnas.org/content/111/2/833.

[55] C. J. Stam et al. “The relation between structural and functional connectivity patterns in complex brain networks”. In: International Journal of Psychophysiology. Research on Brain Oscillations and Connectivity in A New Take-Off State 103 (May 1, 2016), pp. 149–160. ISSN: 0167-8760. DOI: 10.1016/j.ijpsycho.2015.02.011. URL: http://www.sciencedirect.com/science/article/pii/S0167876015000410.

[56] Jil Meier et al. “A Mapping Between Structural and Functional Brain Networks”. In: Brain Connectivity 6.4 (2016), pp. 298–311. DOI: 10.1089/brain.2015.0408. eprint: https://doi.org/10.1089/brain.2015.0408. URL:https://doi.org/10.1089/brain.2015.0408.

[57] Cassiano Becker et al. “Spectral mapping of brain functional connectivity from diffusion imaging”. In: Scientific Reports 8.1411 (2018). DOI: 10.1038/s41598-017-18769-x. URL: https://doi.org/10.1038/s41598-017-18769-x.

[58] Samuel Deslauriers-Gauthier et al. “A unified framework for multimodal structure-function mapping based on eigenmodes”. In: Medical Image Analysis 66 (2020), p. 101799. eprint: doi:10.1016/j.media.2020.101799.

[59] Prejaas Tewarie et al. “Mapping functional brain networks from the structural connectome: Relating the series expansion and eigenmode approaches”. In: Neuroimage 216 (2020), p. 116805. eprint: doi: 10.1016/j.neuroimage.2020.116805.

[60] Jean Daunizeau, Stefan J. Kiebel, and Karl J. Friston. “Dynamic causal modelling of distributed electromagnetic responses”. In: NeuroImage 47.2 (2009). Publisher: Elsevier Inc., pp. 590–601. ISSN: 10538119. DOI: 10.1016/j.neuroimage.2009.04.062. URL:http://dx.doi.org/10.1016/j.neuroimage.2009.04.062.

[61] Klaas E Stephan et al. “Nonlinear dynamic causal models for fMRI”. In: NeuroImage 42.2 (2008), pp. 649–662. DOI: 10.1016/j.neuroimage.2008.04.262.Nonlinear. URL: http://www.ncbi.nlm.nih.gov/pmc/articles/PMC2636907/.

[62] D. A. Pinotsis et al. “Linking canonical microcircuits and neuronal activity: Dynamic causal modelling of laminar recordings”. In: NeuroImage 146 (Feb. 1, 2017), pp. 355–366. ISSN: 1053-8119. DOI: 10.1016/j.neuroimage.2016.11.041. URL:http://www.sciencedirect.com/science/article/pii/S1053811916306590.

[63] Adeel Razi et al. “Construct validation of a DCM for resting state fMRI”. In: NeuroImage 106 (2015). Publisher: The Authors, pp. 1–14. ISSN: 10959572. DOI: 10.1016/j.neuroimage.2014.11.027. URL: http://dx.doi.org/10.1016/j.neuroimage.2014.11.027.

[64] Hae Jeong Park et al. “Dynamic effective connectivity in resting state fMRI”. In: NeuroImage 180 (May 2017 2018). Publisher: The Authors, pp. 594–608. ISSN: 10959572. DOI: 10.1016/j.neuroimage.2017.11.033. URL: https://doi.org/10.1016/j.neuroimage.2017.11.033.

[65] Maria Giulia Preti, Thomas AW Bolton, and Dimitri Van De Ville. “The dynamic functional connectome: State-of-the-art and perspectives”. In: NeuroImage. Functional Architecture of the Brain 160 (Oct. 15, 2017), pp. 41–54. ISSN: 1053-8119. DOI: 10.1016/j.neuroimage.2016.12.061. URL: http://www.sciencedirect.com/science/article/pii/S1053811916307881.

[66] Frederik Van de Steen et al. “Dynamic causal modelling of fluctuating connectivity in resting-state EEG”. In: NeuroImage 189 (Apr. 1, 2019), pp. 476–484. ISSN: 1053-8119. DOI: 10.1016/j.neuroimage.2019.01.055. URL: http://www.sciencedirect.com/science/article/pii/S1053811919300552.

[67] Stefan Frässle et al. “A generative model of whole-brain effective connectivity”. In: NeuroImage 179.May (2018), pp. 505–529. ISSN: 10959572. DOI: 10.1016/j.neuroimage.2018.05.058.

[68] Stefan Frässle et al. “Regression DCM for fMRI”. In: NeuroImage 155 (February 2017). Publisher: Elsevier, pp. 406–421. ISSN: 10959572. DOI: 10.1016/j.neuroimage.2017.02.090. URL: http://dx.doi.org/10.1016/j.neuroimage.2017.02.090.

[69] Adeel Razi et al. “Large-scale DCMs for resting-state fMRI”. In: Network Neuroscience 1.3 (Oct. 27, 2017). Publisher: MIT Press One Rogers Street, Cambridge, MA 02142-1209 USA, pp. 222–241. ISSN: 2472-1751. DOI: 10.1162/NETN_a_00015. URL: https://www.mitpressjournals.org/doi/abs/10.1162/NETN_a_00015.

[70] Daniel S. Margulies et al. “Situating the default-mode network along a principal gradient of macroscale cortical organization”. In: Proceedings of the National Academy of Sciences 113.44 (Nov. 1, 2016). Publisher: National Academy of Sciences Section: Biological Sciences, pp. 12574–12579. ISSN: 0027-8424, 1091-6490. DOI: 10.1073/pnas.1608282113. URL: https://www.pnas.org/content/113/44/12574.

[71] Jorge Sepulcre et al. “Stepwise Connectivity of the Modal Cortex Reveals the Multimodal Organization of the Human Brain”. In: Journal of Neuroscience 32.31 (Aug. 1, 2012). Publisher: Society for Neuroscience Section: Articles, pp. 10649–10661. ISSN: 0270-6474, 1529-2401. DOI: 10.1523/JNEUROSCI.0759-12.2012. URL: https://www.jneurosci.org/content/32/31/10649.

[72] Bertha Vázquez-Rodríguez et al. “Gradients of structure–function tethering across neocortex”. In: Proceedings of the National Academy of Sciences (Sept. 30, 2019). Publisher: National Academy of Sciences, p. 201903403. ISSN: 0027-8424. DOI: 10.1073/PNAS.1903403116. URL: https://www.pnas.org/content/early/2019/09/27/1903403116 (visited on 09/30/2019).

[73] Julia M. Huntenburg, Pierre-Louis Bazin, and Daniel S. Margulies. “Large-Scale Gradients in Human Cortical Organization”. In: Trends in Cognitive Sciences 22.1 (Jan. 1, 2018), pp. 21–31. ISSN: 1364-6613. DOI: 10.1016/j.tics.2017.11.002. URL: http://www.sciencedirect.com/science/article/pii/S1364661317302401 (visited on 08/26/2020).

[74] Randy L. Buckner and Daniel S. Margulies. “Macroscale cortical organization and a default-like apex transmodal network in the marmoset monkey”. In: Nature Communications 10.1 (Apr. 29, 2019). Number: 1 Publisher: Nature Publishing Group, p. 1976. ISSN: 2041-1723. DOI: 10.1038/s41467-019-09812-8. URL: https://www.nature.com/articles/s41467-019-09812-8 (visited on 08/26/2020).

[75] William W. Seeley et al. “Dissociable Intrinsic Connectivity Networks for Salience Processing and Executive Control”. In: Journal of Neuroscience 27.9 (Feb. 28, 2007). Publisher: Society for Neuroscience Section: Articles, pp. 2349–2356. ISSN: 0270-6474, 1529-2401. DOI: 10.1523/JNEUROSCI.5587-06.2007. URL: https://www.jneurosci.org/content/27/9/2349.

[76] Michael D. Fox et al. “The human brain is intrinsically organized into dynamic, anticorrelated functional networks”. In: Proceedings of the National Academy of Sciences 102.27 (July 5, 2005). Publisher: National Academy of Sciences Section: Biological Sciences, pp. 9673–9678. ISSN: 0027-8424, 1091-6490. DOI: 10.1073/pnas.0504136102. URL:https://www.pnas.org/content/102/27/9673.

[77] Bruce Fischl et al. “Whole Brain Segmentation: Automated Labeling of Neuroanatomical Structures in the Human Brain”. In: Neuron 33 (2002), pp. 341–355.

[78] Jennifer A. McNab et al. “The Human Connectome Project and beyond: Initial applications of 300mT/m gradients”. In: NeuroImage (2013). ISSN: 10538119. DOI: 10.1016/j.neuroimage.2013.05.074.

[79] Mark Jenkinson et al. “FSL”. In: NeuroImage 62.2 (Aug. 2012), pp. 782–790. ISSN: 1053-8119. DOI: 10.1016/J.NEUROIMAGE.2011.09.015. URL:https://www.sciencedirect.com/science/article/pii/S1053811911010603?via%7B%5C%%7D3Dihub.

[80] Bruce Fischl. “FreeSurfer”. In: NeuroImage 62.2 (Aug. 2012), pp. 774–781. ISSN: 10538119. DOI: 10.1016/j.neuroimage.2012.01.021. URL: http://www.ncbi.nlm.nih.gov/pubmed/22248573 http://www.pubmedcentral.nih.gov/articlerender.fcgi?artid=PMC3685476 https://linkinghub.elsevier.com/retrieve/pii/S1053811912000389.

[81] David J. Wales and Jonathan P.K. Doye. “Global optimization by basin-hopping and the lowest energy structures of Lennard-Jones clusters containing up to 110 atoms”. In: Journal of Physical Chemistry A 101.28 (1997), pp. 5111–5116. ISSN: 10895639. DOI: 10.1021/jp970984n. URL: https://pubs.acs.org/doi/10.1021/jp970984n.

[82] Xihe Xie, Megan J. Stanley, and Pablo F. Damasceno. “Raj-Lab-UCSF/spectrome: Spectral Graph Model of Connectomes (Version 0.15)”. In: zenodo (Nov. 6, 2019). DOI: 10.5281/ZENODO.3532497. URL: https://zenodo.org/record/3532497 (visitedon 11/07/2019).

